# Functional similarity, not phylogenetic relatedness, predicts the relative strength of competition

**DOI:** 10.1101/2021.07.21.453226

**Authors:** Michael B. Mahon, David E. Jennings, David J. Civitello, Marc J. Lajeunesse, Jason R. Rohr

## Abstract

Predicting the outcome and strength of species interactions is a central goal of community ecology. Researchers have proposed that outcomes of species interactions (competitive exclusion and coexistence) are a function of both phylogenetic relatedness and functional similarity. Studies relating phylogenetic distance to competition strength have shown conflicting results. Work investigating the role of phylogenetic relatedness and functional similarity in driving competitive outcomes has been limited in terms of the breadth of taxa and ecological contexts examined, which makes the generality of these studies unclear. Consequently, we gathered 1,748 pairwise competition effect sizes from 269 species and 424 unique species pairs with divergence times ranging from 1.14 to 1,275 million years and used meta-regression and model selection approaches to investigate the importance of phylogenetic relatedness and functional similarity to competition across ecological contexts. We revealed that functional similarity, but not phylogenetic relatedness, predicted the relative strength of interspecific competition (defined as the strength of interspecific competition relative to intraspecific competition). Further, we found that the presence of predators, certain habitats, increasing density of competitors, and decreasing spatial grain of experiments were all associated with more intense interspecific competition relative to intraspecific competition. Our results demonstrate that functional similarity, not phylogenetic relatedness, may explain patterns of competition-associated community assembly, highlighting the value of trait-based approaches in clarifying biotic assembly dynamics.

## Introduction

Predicting the outcomes of species interactions and community assembly and disassembly processes have long been the foci of community ecologists^1^. A core prediction of community ecology, originally described by Darwin^2^, is that closely related species will compete more strongly with each other than distantly related species, referred to as the “competition-relatedness hypothesis”^3^. Due in part to advances in genetics, molecular biology, and phylogenetic tools in the last 25 years, Darwin’s hypothesis has undergone repeated empirical tests. However, the generality of the relationship between phylogenetic relatedness and competition strength remains uncertain, as studies show conflicting results^3–12^.

Conflicting empirical patterns between phylogeny and competition may suggest that the assumptions underlying Darwin’s hypothesis may be flawed^7^. Darwin’s hypothesis assumes that phylogenetically related species are more ecologically similar than distantly related species (i.e., phylogenetic niche conservatism)^2–4,13,14^. While some support exists for phylogenetic niche conservatism^4,15–17^, niches can evolve convergently or randomly^18–20^, suggesting that phylogenetically similar species may not be ecologically similar^21^. Darwin’s hypothesis also assumes that ecologically similar species compete more strongly than less similar species, because they share similar niches^2–4,13,14^. However, niche differences (i.e., degree of niche overlap) alone may not predict competition strength; rather, competitive ability differences (i.e., differences in species’ abilities to utilize shared limiting resources) may drive patterns of competitive exclusion, while niche overlap may determine whether species pairs can coexist^22–24^. This suggests that ecologically similar species may not inherently compete more strongly than ecologically dissimilar species. Specifically, coexistence theory predicts competitive exclusion when species with overlapping niches differ in their competitive abilities, and coexistence when strong niche differentiation overcomes differences in competitive ability between species^22–24^. Thus, without information on both the niche and competitive ability of interacting species, successfully predicting the strength of competition remains challenging.

Experimental efforts to quantify factors that influence competition strength have been limited by the breadth of focal taxa^25^ and/or the experimental and ecological contexts of observations^26,27^. For instance, competition strength has been shown to be higher in certain taxonomic groups and habitat types^25,28^. Additionally, certain experimental conditions may intensify competition between species; specifically, studies have shown that competition is stronger when predators are present^26,29^, under resource limitation^30,31^, or when experiments are conducted in mesocosms rather than natural field settings^27,32^. Thus, certain experimental conditions might obscure patterns among phylogenetic relatedness, functional similarity, and competition strength.

We propose to address a central goal in ecology by resolving the uncertainty regarding the influences of phylogenetic relatedness and functional similarity on competition. Resolving this uncertainty will improve predictions concerning the outcome of species interactions and increase understanding of biotic mechanisms involved in the assembly and structuring of communities^33–37^. Here, we use a meta-analysis of 1,748 effect sizes of pairwise competitive interactions across a range of taxa and ecological contexts to assess whether the strength of interspecific competition relative to the strength of intraspecific competition (referred to here as competitive response^38,39^) is better predicted by phylogenetic relatedness or functional similarity of competing species. Competitive response represents the ability of a species to tolerate competition from other species^38^ and translates to the long-term winners and losers of competition (i.e., a species that experiences low interspecific competition relative to intraspecific competition will eventually competitively exclude a competing species that is less tolerant of interspecific competition)^39^. Our objectives are to: 1) investigate the relationships among phylogenetic distance (time since divergence), functional similarity (body size of focal species relative to body size of competing species), and competitive response across taxonomic groups and 2) determine the ecological and experimental contexts that may influence the response to competition. Per Darwin’s competition-relatedness hypothesis^2^, we predict that competitive response is negatively related to phylogenetic distance (e.g., stronger relative interspecific competition in closely related species pairs) and, per coexistence theory^23^, we predict that competitive response is positively related to functional similarity (e.g., stronger relative interspecific competition when the focal species is smaller than the competing species; size-asymmetric competition^40^). Further, we hypothesize that certain ecological and experimental contexts will influence a species’ response to competition. For instance, we predict that 1) resource limitation, 2) relatively high densities of the competing species, 3) the presence of predators, and 4) interactions in closed experimental systems compared to those in open, field systems will increase interspecific competition relative to intraspecific competition.

## METHODS

We used Web of Science to search eight ecology journals (*American Naturalist*, *Ecological Monographs*, *Ecology*, *Journal of Animal Ecology*, *Journal of Ecology*, *Journal of Experimental Marine Biology and Ecology*, *Oecologia*, and *Oikos*) from 1988-2008 for the keywords ‘interspecific competition’ on 20 June 2008, which yielded 1,039 studies. A second search for the same keywords within the same subset of ecological journals from 2008-2020 was conducted on 3 March 2021, which yielded an additional 399 studies. In total, our searches yielded 1,438 studies. After studies were collected, they were examined to determine if they met seven criteria for inclusion in our meta-analyses: 1) the study species had phylogenetic information (at the species or genus level) deposited in the TimeTree database^41,42^, an online comprehensive list of estimated divergence times across all the major taxonomic groups, 2) the density of at least one of the study species was manipulated experimentally, 3) the experimental design used appropriate controls (e.g., non-manipulated groups of individuals), 4) the means, variance, and sample sizes were reported or displayed (data from figures were extracted using the imageJ software^43^) or data were available, allowing us to calculate effect sizes (studies with single replicates were excluded from the analyses, because effect sizes could not be reliably calculated from these data), 5) clear pair-wise interactions were present in the study (e.g., species A and species B were grown together and species A was grown separately), 6) the endpoint measured was broadly applicable across different taxonomic groups (e.g., a measure of biomass, density, or survival, as opposed to a taxon-specific variable such as time to metamorphosis), and 7) the species studied had published information on mean size at maturity available from the study itself, the USDA Plants Database (www.plants.usda.gov), or Animal Diversity Web (https://animaldiversity.org/). In total, 96 studies with 1,748 effect sizes met our inclusion criteria.

Once studies were determined to meet the criteria for inclusion, we collected data on means, variances, sample sizes, the number of individuals of the species pairs, habitat (aquatic or terrestrial), whether the experiment also manipulated predator (or herbivore) presence and/or resource levels, the experimental venue (field or mesocosm), the spatial grain of the experiment (area or volume of study; m^2^ or liters), and the length of experiment (days). A mesocosm venue was defined as any lab, greenhouse, or outdoor experiment conducted in a bounded and partially enclosed venue (e.g., beaker in the lab, pots in a greenhouse, outdoor aquatic tanks), and a field venue was defined as any experiment conducted in a natural setting outside of an enclosed venue (e.g., managed field system, streams, forests). We calculated relative spatial grain as the log10 transformation of spatial grain divided by body size of the focal species (species A).

Phylogenetic relatedness was calculated as the log10 transformation of time since the divergence (mya) of the competing species (median value from TimeTree^41,42^ recorded on 20 March 2021). We calculated relative density as the log10 transformation of ratio of the number of individuals of the focal species (species A) to the number of individuals of the competing species (species B). Functional similarity was estimated as the log10 transformation of the ratio of body size (cm) at maturity of the focal species (species A) to the body size of the competing species (species B; relative body size).

### Effect sizes

We calculated the effect sizes for the outcome of species interactions using the Delta-method-adjusted LRR (LRR^Δ^; equation (1)), as log response ratio (LRR) can be biased when estimating the outcome of studies with small sample sizes^44^.

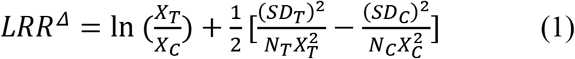

Where *X_T_* is the mean of the treatment group (focal species grown with heterospecifics), *X_C_* is the mean of the control group (focal species grown with conspecifics), and SD and N are the within-study standard deviations and sample sizes, respectively, of the treatment and control groups. Effective sample sizes (N) were calculated as the number of replicates multiplied by the number of individuals per replicate. Here, LRR^Δ^ is the ratio between interspecific and intraspecific effects and can be interpreted as the competitive response of a species (ability of a species to avoid being suppressed; *sensu* refs^38,39^). Specifically, when LRR^Δ^ is negative, the focal species is more responsive to interspecific effects than intraspecific effects, indicating that the species is sensitive to interspecific competition and long-term competitive exclusion of the focal species can occur^39^. Conversely, when LRR^Δ^ is zero or positive, interspecific effects are equal to or less than intraspecific effects, indicating that the species is not sensitive to interspecific competition and the focal species may, in the long-term, competitively exclude the competing species^39^. As some effect sizes within studies were calculated using the same control group and their observations were therefore non-independent (equation (2)), we calculated variance-covariance matrices for observations (**A** and **B** below) within studies that shared control groups using the ‘covariance_commonControl’ function in the ‘metagear’ R package^45^ and adjusted per the Delta-method^44^.

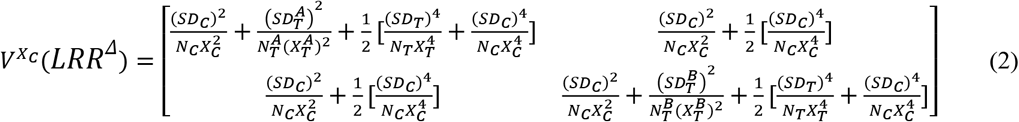

### Statistical analyses

All analyses were conducted using R 4.0.3^46^. We used the package ‘metafor’^47^ and the ‘rma.mv’ function to generate mixed effects meta-regression models, described with equation (3).

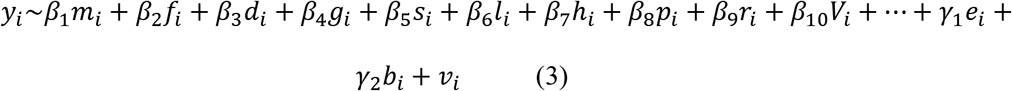

Where *y_i_* denotes the Delta-adjusted LRR and *v_i_* denotes the Delta-adjusted LRR variance for the *i*th effect size. Our effect sizes have a hierarchical structure; multiple effect sizes exist within single experiments. To minimize the risk of Type I error associated with the nonindependence among effect sizes that were not explained by sampling error alone^48,49^, we accounted for this nonindependence by 1) including a random effects component for effect sizes within studies (*e*) and between studies (*b*) and 2) estimating sampling covariance among effect sizes that have shared control groups (equation (2); *v*). We did not assess potential publication bias, because the nonindependence among effect sizes within a study and the resulting variance-covariance matrix invalidates these tests^48,50^.

Moderators in the mixed effects meta-analytic models included log10 phylogenetic distance (*m*), log10 relative body size (*f*), log10 relative density (*d*), log10 relative spatial grain (*g*), study endpoint (density/growth/survival; *s*), log10 length of experiment (days; *l*) habitat (terrestrial/aquatic; *h*), predators (present/absent; *p*), resource level (low/ambient/high; *r*), venue (field/mesocosm; *V*), and all possible two-way interactions (… in equation (3); interactions between both resource and study endpoint and any other categorical moderator as well as between venue and habitat were not possible due to missing cells or highly unbalanced replication across cells). To evaluate the importance of these moderators, we performed model selection based on AIC in which we fit all possible combinations of main effects and two-way interactions as moderators using the ‘dredge’ function in the ‘MuMIn’ R package^51^. To prevent overfitting of models, we limited models to 8 moderators (main and interactive effects). Model weights (AIC*w*) and relative importance values (sum of AIC*w* across models in which the focal moderator appears) for each moderator were calculated from models with a ΔAIC ≤ 4 (see Supp. Table S1 and Supp. Fig. S1 for model weights and moderator importance values)^52^. To test whether continuous moderators were different from 0, we used z-tests. Similarly, to test whether categorical moderators were different from 0, we used z-tests and pairwise differences among groups within categorical moderators were evaluated with t-test contrasts. To estimate and plot marginal effects of moderators, we calculated estimated marginal means; continuous covariate moderators were held at their median values and categorical covariate moderators were averaged over the proportions of the groups. Summary statistics, p-values, and figures were generated from the model with the lowest AIC.

Following the primary analyses described above, we examined the effect of resource limitation on endpoints for species grown with conspecific and heterospecific neighbors, by selecting the 14 studies in our database that manipulated resources (e.g., ambient and high, ambient and low, or low and high resources). In each study, for all species that were grown with both conspecific and heterospecific neighbors, we calculated the Delta-method-adjusted LRR (LRR^Δ^; equation (1)) with “treatment” set as resource limitation and “control” set as higher resource conditions for each species-neighbor combination, which resulted in 150 effect sizes. As with the primary analyses, we calculated a variance-covariance matrix from equation (2) and used the ‘rma.mv’ function to generate a mixed effects meta-analysis model with random effects components for effect sizes within studies and between studies. To test whether resource limitation differentially affected inter- and intra-specific competition, we included a single moderator: the identity of the neighbor(s) of the focal species (conspecific/heterospecific).

## RESULTS

We collected 1,748 effect sizes from 96 studies (involving 269 different species and 424 species pairs) that met our inclusion criteria. The divergence times of the species obtained from the TimeTree database ranged from 1.14 million years ago (MYA) to 1,275 MYA, with a median divergence time of 121 MYA. Species belonged to various taxonomic groups (49.3% angiosperms, 17.2% amphibians, 11.8% arthropods, 8.1% fish, 4.1% mollusks, and 3.2% mammals, with the remainder from bryophytes, echinoderms, gymnosperms, reptiles, and rotifers), and data represent different measures of competition (88.7% growth, 8.9 % survival, and 2.4% density). Effect sizes were distributed among various ecological contexts (Fig. 1) and were not different among broad taxonomic groupings of focal species (Supp. Fig. S2) or among study endpoints (Supp. Fig. S3).

**Figure 1.**
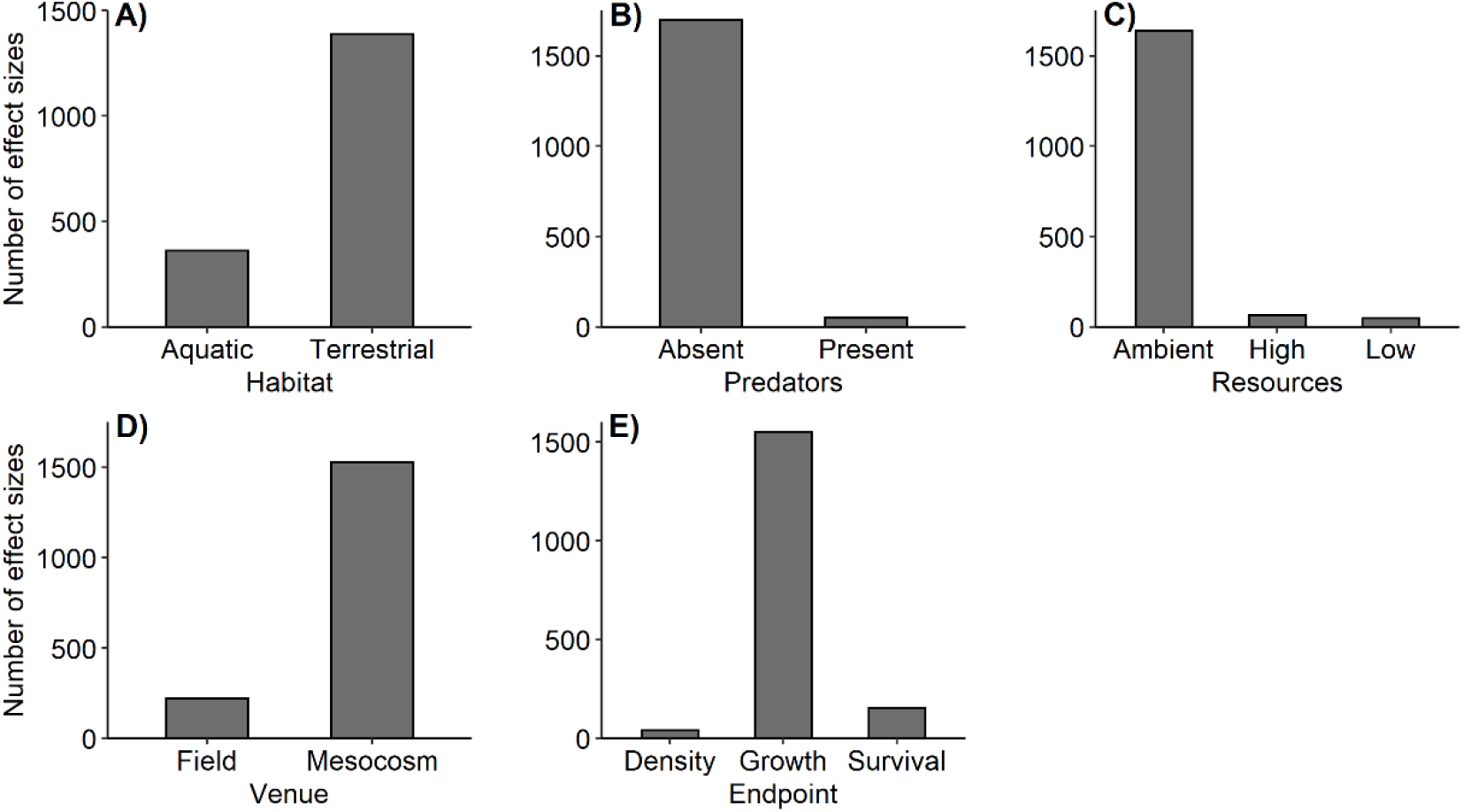
Representation of effect size data for different ecological contexts. Shown are a) habitat (aquatic or terrestrial), b) whether the experiment manipulated the predator presence, c) whether the experiment manipulated resource levels, d) the experimental venue (field or mesocosm), and e) the endpoint measured.

When controlling for within- and among-study variance, model selection indicated that competitive response was negatively related to relative body size of the focal species (z = 3.944, p < 0.001; Table 1, Fig. 2A, Supp. Fig. S4). Specifically, when the focal species was smaller than the competing species, interspecific competition was greater than intraspecific competition, but interspecific competition was less than intraspecific competition when individuals of the focal species were larger than the competing species. The magnitude of this effect increased when predators were present (z = 2.682, p = 0.007; Table 1, Fig. 2A, Supp. Fig. S4).

**Figure 2.**
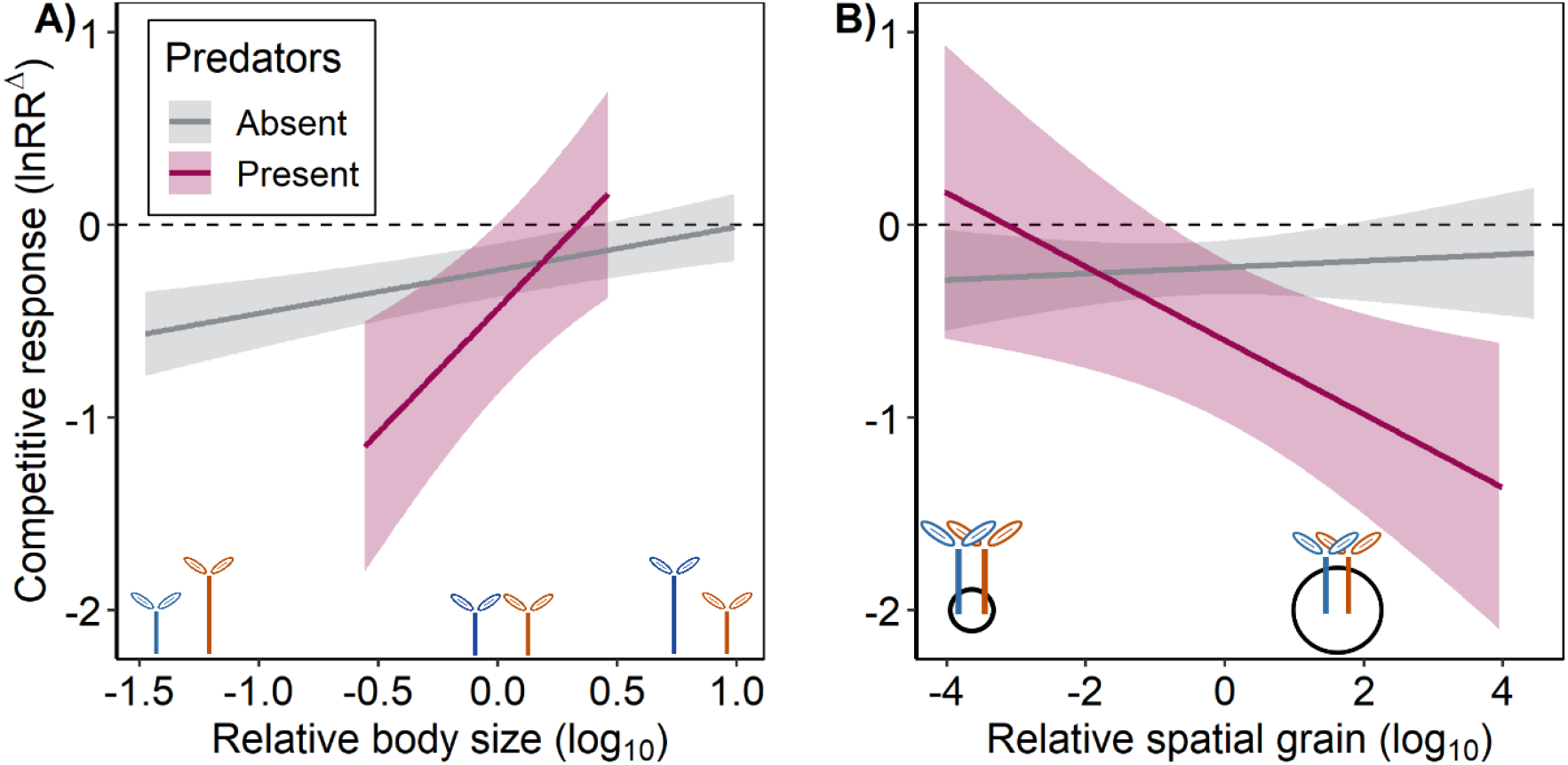
Marginal effects plot showing the interactive effects of predation and (A) relative body size (size of focal species (blue) / size of competing species (orange); functional similarity) and (B) relative spatial grain (spatial grain / size of focal species; size of black circle relative to blue individual) on the outcome of species interactions (lnRR^Δ^), respectively. In (A), competition strength decreases as the focal species (Sp1, blue) increased in size relative to the competing species (Sp2, orange), suggesting that relative functional differences drive competition strength. The magnitude of this effect is increased in the presence of predators. In (B), the presence of predators increases competition strength with relative spatial grain is high, but not when relative spatial grain is low, suggesting that the presence of predators may alter spatial resource use by competing species (increased aggregation) in experiments with relatively large spatial grain. Lines represent marginal effects of (A) relative body size and (B) relative spatial grain included in the mixed-effects meta-regression model and shading shows associated 95% credible bands. To estimate these marginal effects, categorical covariates from the mixed-effects meta-regression model were held at their proportional values and continuous covariates were held at their median value. Model-predicted regression lines and credible bands are shown with data points in Fig. S4 and S5 in the supplement.

**Table 1:** Results of a mixed-effects meta-regression model from the model selection indicated best model (lowest AIC) relating competition strength to factors including the relative size of the focal species, relative density of the focal species, presence of predators, and habitat. Variance estimates of the within and across study random effects are 0.390 and 0.257, respectively.

| Moderators | Coefficient | 95% CI | Z-value | P-value |
| --- | --- | --- | --- | --- |
| Grand Mean | -0.239 | 0.18 | -2.607 | **0.009** |
| Relative Body Size (log10) † | 0.224 | 0.11 | 3.944 | **<0.001** |
| Relative Density (log10) † | 0.234 | 0.13 | 3.510 | **<0.001** |
| Relative Spatial Grain (log10) † | 0.017 | 0.06 | 0.512 | **0.609** |
| Predators Present | 0.983 | 0.44 | 4.387 | **<0.001** |
| Predators Present * Relative Body Size | 1.062 | 0.78 | 2.682 | **0.007** |
| Predators Present * Relative Spatial Grain | -0.208 | 0.15 | -2.656 | **0.008** |
| Terrestrial Habitat | 0.024 | 0.26 | 0.181 | 0.856 |
| Predators Present * Terrestrial Habitat | -1.715 | 0.72 | -4.687 | **<0.001** |

Notes: The average outcome of competition is indicated by the grand mean, as the coefficient represents the pooled outcome of competition. For continuous moderators (†), bolded p-values indicate a slope that deviates significantly from zero and the sign of the coefficient indicates the direction of the effect. For categorical moderators, bolded p-values indicate a significant difference from the grand mean with the coefficient indicating the direction and the estimated mean of the category. For interactions between categorical and continuous moderators, bolded p-values indicate a slope that deviates significantly from the continuous moderator and the sign of the coefficient indicates the direction of the effect.

The presence of predators also interacted with several ecological moderators to affect the competitive response of species (Table 1). Specifically, there was an interaction between relative spatial grain of the study and the presence of predators, such that when predators were absent, there was a weak positive relationship between relative spatial grain and competitive response (β: 0.017; Table 1, Fig. 2B, Supp. Fig. S5), but when predators were present, competitive response was negatively related to relative spatial grain (β: −0.191; Table 1, Fig. 2B, Supp. Fig. S5). Further, predator presence interacted with habitat to affect the competitive response of species (z = −3.459, p = <0.001). In terrestrial habitats, the presence of predators increased competitive response (marginal lnRR^Δ^ = −0.78 ± 0.53 95% CI) relative to when predators were absent (marginal lnRR^Δ^ = −0.23 ± 0.17 95% CI; pairwise comparison, t = 1.966, p = 0.049; Fig. 3). Conversely, in aquatic systems, the presence of predators reduced competitive response (marginal lnRR^Δ^ = 0.91 ± 0.56 95% CI) relative to when predators were absent (marginal lnRR^Δ^ = −0.25 ± 0.21 95% CI; pairwise comparison, t = −3.799, p < 0.001; Fig. 3).

**Figure 3.**
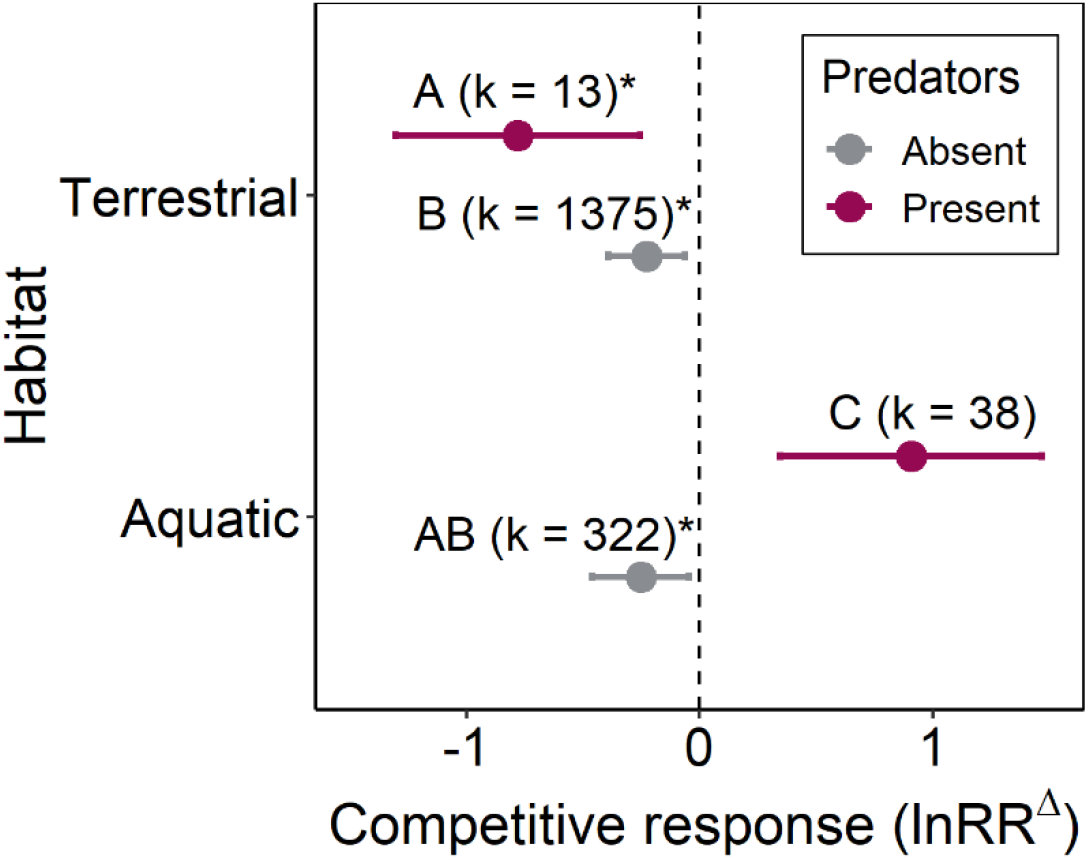
Interaction plot showing the marginal interactive effect of habitat (aquatic or terrestrial) and predators (absent or present) on competitive response. Pooled effects and errorbars (95% CI) were generated from the mixed-effects meta-regression model (see Table 1). The coloration of points and errorbars corresponds with predators (grey are predators absent and purple is predators present). Factor combinations sharing the same letter are not different from each other in pairwise comparisons. Asterisks represent factor combination estimates that are significantly different from 0. Above each estimate are counts of effect sizes for each factor combination. There are no differences in competition strength between aquatic and terrestrial habitats when predators are absent. When predators are present, competition strength increases in terrestrial habitats, but decreases in aquatic habitats. To estimate these pooled effects, continuous covariates from the mixed-effects meta-regression model were held at their median values.

Finally, model selection also indicated that the relative density of the focal species to the competing species was negatively related to competitive response (z = 3.510, p < 0.001; Table 1, Fig. 4, Supp. Fig. S6). Specifically, when densities of the focal species were less than the densities of the competing species, interspecific competition was greater than intraspecific competition, whereas the opposite was true when the densities of the focal species were greater than the densities of the competing species. Despite functional similarity and phylogenetic relatedness being positively correlated (simple linear regression: F_1,422_ = 11.59, p < 0.001, R^2^ = 0.03; Supp. Fig. S7), we found no evidence that phylogenetic distance alone or when controlling for functional distance was a significant predictor of competitive response (Supp. Tables S1, S2; Supp. Fig. S1). Additionally, analysis of effect sizes comparing resource limitation on inter- and intraspecific competition indicated that resource limitation consistently intensifies competition, regardless of whether focal species were with hetero- or conspecifics (effect of resource limitation on intra- vs. inter-specific competition, Z = −0.01, p = 0.88; Supp. Fig. S8). Thus, resource limitation does not appear to alter the ratio of inter-to intraspecific competition (competitive response; our measure of competition strength). Rather, on average, it seems to equally intensify inter- and intraspecific competition. Finally, we found little evidence for study endpoint, study duration, or venue influencing competitive response (Supp. Table S1, Supp. Fig. S1).

**Figure 4.**
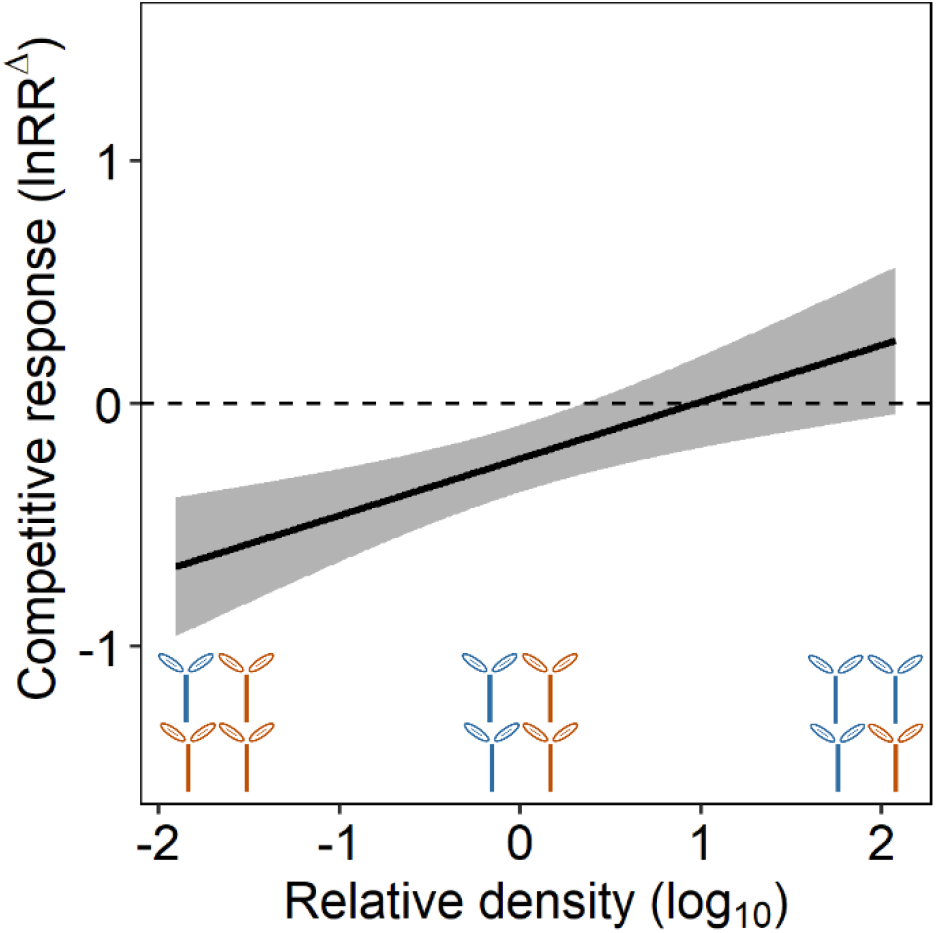
Marginal effects plot showing the effects of relative density (density of focal species (blue)/ density of competing species (orange)) on the outcome of species interactions (lnRR^Δ^). Competition strength decreases as the focal species (Sp1, blue) increased in density relative to the competing species (Sp2, orange), suggesting that relative densities of competing species can influence the outcome of the interaction and, thus, potentially obscure generalities from individual competition studies. Lines represent marginal effects of relative density included in the mixed-effects meta-regression model and shading shows associated 95% credible bands. To estimate these marginal effects, categorical covariates from the mixed-effects meta-regression model were held at their proportional values and continuous covariates were held at their median value. Model-predicted regression line and credible band are shown with data points in Fig. S6 in the supplement.

## DISCUSSION

Predicting the outcome of competition, and thus, the mechanisms underlying species coexistence patterns and biotic assembly processes within communities, is a core tenet of community ecology^1^. A long held assumption was that the strength of interspecific competition is driven by phylogenetic relatedness, with closely related species competing more strongly than distantly related species^2–4^. Previous investigations of phylogenetic relatedness and functional similarity effects on competition strength have been limited by the breadth of focal taxa examined^3,7,25^ and/or the experimental and ecological contexts in which observations occurred^26,27^. Using a meta-analysis of 1,748 effect sizes across a range of taxa and ecological contexts, we evaluated whether phylogenetic relatedness and/or functional similarity explained species’ competitive responses and whether specific ecological contexts influenced the competitive response of species. Our results suggest that functional similarity, and not phylogenetic relatedness, predicts the competitive response of species (i.e., strength of interspecific competition relative to strength of intraspecific competition). Further, we found that certain ecological and experimental contexts, such as habitat, the presence of predators, relative spatial grain, and relative densities of competing species, can directly alter a species’ competitive response.

Our findings add to an increasing number of studies that do not support Darwin’s hypothesis that phylogenetic relatedness drives competition strength^4,5,7,53–55^, suggesting that one or both underlying assumptions of this hypothesis are invalid^7^. Examples of phylogenetic niche conservatism exist^4,15–17^, but niches can evolve convergently or randomly^18–20^, suggesting that phylogenetic and ecological relatedness may not always be correlated. Regardless of whether niche overlap between species pairs is related to their phylogenetic relatedness, the degree of niche overlap might not exclusively drive competition; rather, competitive ability differences (i.e., differences in species’ abilities to utilize shared limiting resources) in addition to niche overlap are thought to influence the outcome of species interactions^22–24^. While there was some evidence for a phylogenetic signal in the functional similarity of species, phylogenetic relatedness was never a predictor of competition strength in our analyses. Therefore, our findings, in conjunction with previous research^6^, suggest that phylogeny is not an effective predictor of the outcome of species interactions.

Our meta-analysis indicated that a species’ competitive response is driven by functional similarity, which may be correlated with differences in competitive ability between species pairs^53,56–58^. Thus, we can infer that a species’ competitive response (sensitivity to interspecific competition) is likely driven by that species’ relative functional ability (e.g., the height of a plant species relative to its competitor), lending further credence to modern coexistence theory^22–24^, which, in contrast to Darwin’s competition-relatedness hypothesis, posits that functional similarity drives competitive exclusion when species pairs have overlapping niches, while niche differences facilitate coexistence. While we did not explicitly test for niche differences between competing species^23^, species pairs within our meta-analysis were those that were likely to compete (i.e., researchers would not select species that do not compete with one another if they were interested in studying competition) and, as such, likely have some niche overlap. Therefore, to thoroughly test modern coexistence theory^22–24^, future research should explicitly test competitive response in an assortment of taxa across a range of both competitive ability and niche differences^32^.

While previous research suggests that certain ecological contexts can influence the effects of phylogenetic relatedness and functional similarity on competition strength^3,4,6,9^, we found that only the presence of predators altered the relationship between functional similarity and competitive response. Specifically, the presence of predators intensified asymmetrical competition^40^, such that the relative strength of interspecific competition was more responsive to the relative size differences between competing species when predators were present than when predators were absent. Aside from predators, no other ecological or experimental factors influenced the relationship between functional similarity and competitive response. Nonetheless, we found evidence that certain ecological and experimental contexts may directly alter the competitive response of species.

Lending support to previous research showing effects of predation on competition^26,29^, we found evidence that the presence of predators, interacting with habitat, can influence competition strength. The interaction between predators and habitat observed here might be related to habitat productivity. Specifically, predators are hypothesized to increase competition in low productivity systems, because 1) predators reduce the range of conditions that allow for coexistence, 2) species that are well-defended from predation are likely poor at attaining resources and are likely to be outcompeted, and/or 3) traits that increase competitive ability may simultaneously increase vulnerability to predation, and, thus, these traits may be less pronounced in low productivity systems relative to high productivity systems^29,59^. Consistent with our results, predation pressures in terrestrial systems, which may be less productive than aquatic systems^60,61^, may increase interspecific competition strength, whereas predation pressures in aquatic systems may reduce or not affect the strength of interspecific competition^29,59^.

Further, we found an interaction between predators and relative spatial grain. When predators were present, but not when they were absent, competitive response was negatively related to relative spatial grain (i.e., interspecific competition was stronger than intraspecific competition in experiments with relatively larger spatial grain), suggesting that spatial refugia are a more limiting resource for the competing species when predators are present than absent^29^. The predator-mediated reduction of space likely increases interactions between competing species^62^, which, in turn, likely leads to increased interspecific competition^26,29^ because species pairs were likely to have some degree of niche overlap.

As predicted, we found a relationship between competitive response and the relative density of the focal species, providing support for density-dependent competition^32,63^. Interspecific competition was greater than intraspecific competition across all taxa when the focal and competing species were at the same densities, suggesting that per capita interspecific competition (e.g., α*_ij_* in Lotka-Volterra competition models) is generally greater than per capita intraspecific competition (e.g., α_ii_ in Lotka-Volterra competition models), in contrast to results found within plant communities^32^. Differences between our results and previous research^32^ may be a product of our inclusion of animal, plant, and cross-kingdom species pairs, as opposed to only plant species pairs, and/or not explicitly estimating the Lotka-Volterra competition coefficients for these competing species^32^. These results also imply that total intraspecific competition pressures may be relatively strong at higher conspecific densities, providing further evidence that the density-dependent mortality observed with Janzen-Connell effects (i.e., greater mortality when conspecifics are aggregated in close proximity) may be, in part, the result of strong intraspecific competition, not just from natural enemies^32,64,65^. To robustly test Janzen-Connell effects, future research should attribute density-dependent mortality to both enemies and intraspecific competition^64^. As we found that certain experimental and ecological factors alter the relative strengths of inter- and intra-specific competition, some discrepancies across studies investigating phylogenetic relatedness and functional similarity controls on competition might be the result of differences in predator presence, habitat, relative size of the experimental venue, and relative densities of competing species. Therefore, future research must account for the influence of these experimental and ecological factors on species’ competitive responses.

While previous research has highlighted the importance of resource availability in driving competition strength^30^, we found no evidence that resource availability affected species’ competitive responses. Rather, we found that resource limitation consistently negatively affected species endpoints, regardless of whether their competitors were hetero- or conspecifics. This result may suggest that resource limitation increases both inter- and intra-specific competition similarly, supporting previous research showing that competition strength increases as resources become limited^30,31^. Alternatively, this result may indicate that resource limitation reduces species performance, but because resource limitation did not affect the ratio between inter- and intraspecific competition, competition strength may be independent of resource availability^66,67^. To clarify the effects of resource availability on competition intensity, future research should explicitly test the strength of inter- and intraspecific competition across a range of taxa and resource levels.

The ability for phylogeny to predict the outcomes of species interactions and biotic assembly processes has been highly debated, with recent evidence suggesting that phylogenetic relatedness alone cannot predict patterns of species coexistence^1,6,7,21,37^. Here, using a meta-analysis of competitive responses across taxa and ecological contexts, we found that functional similarity, a proxy for differences in competitive ability, but not phylogenetic relatedness, was key to predicting species responses to competition, lending support for coexistence theory^22,23^. The lack of a phylogenetic signal in competitive response across broad taxonomic and ecological contexts found here, in conjunction with previous studies^3,6,7,9^, provides considerable evidence against Darwin’s competition-relatedness hypothesis and, thus, against the hypothesis that phylogeny predicts the outcome of species interactions. Our results demonstrate that functional similarity may explain patterns of competition-associated community assembly, thereby highlighting the value of trait-based approaches in clarifying biotic assembly dynamics.

## Author contributions

MBM, DEJ and JRR conceived the study; DEJ and MBM conducted literature searches; DEJ, MBM, DJC, and MJL contributed to the analyses; MBM, DEJ, and JRR wrote the manuscript; all authors provided comments on the manuscript.

## Data and code sharing and accessibility

All data and code will be made available via Dryad and Github, respectively, following acceptance.

## ACKNOWLEDGEMENTS

This study was supported by a Fern Garden Club Scholarship and Rutlish Foundation Grant to D.E.J., and a University of South Florida New Researcher Grant (RO65462) to J.R.R. We thank members of the Rohr lab for providing helpful comments for improving the manuscript.

