## Supplementary material for "Functional similarity, not phylogenetic relatedness, predicts the relative strength of competition": Supp.

**Supplemental Materials**

Supplemental Table 1. Best and competing models (ΔAIC < 4) for competition strength (results presented in main text). Moderators were habitat, venue, predator presence, relative spatial grain (Rel. Spatial Grain), resources, study endpoint, relative density (Rel. Density), phylogenetic distance, and relative body size (Rel. Size). Model selection allowed for the main effect of all moderators, two-way interactions between all factors (interaction between 'predators' and 'model endpoint' was excluded because of lack of replication). Models were mixed-effects meta-regression models that included a random effect of 'study' and had a variance-covariance matrix as described in the main text.

| Moderators | df | logLik | AIC | ΔAIC | AIC*w* |
|---|---|---|---|---|---|
| Rel. Density, Habitat, Rel. Spatial Grain, Predators, Rel. Size, Predators:Habitat, Predators:Rel. Spatial Grain, Predators:Rel. Size | 11 | -1908.90 | 3839.81 | 0.00 | 0.52 |
| Habitat, Rel. Spatial Grain, Predators, Rel. Size, Predators:Habitat, Habitat:Rel. Size, Predators:Rel. Spatial Grain, Predators:Rel. Size | 11 | -1910.20 | 3842.39 | 2.59 | 0.14 |
| Rel. Density, Habitat, Predators, Rel. Size, Venue, Predators:Habitat, Predators:Rel. Size, Predators:Venue | 11 | -1910.47 | 3842.95 | 3.14 | 0.11 |
| Rel. Density, Habitat, Predators, Rel. Size, Predators:Habitat, Predators:Rel. Size | 9 | -1912.74 | 3843.49 | 3.68 | 0.08 |
| Rel. Density, Habitat, Predators, Rel. Size, Predators:Habitat, Habitat:Rel. Size, Predators:Rel. Size | 10 | -1911.82 | 3843.65 | 3.84 | 0.08 |
| Habitat, Rel. Spatial Grain, Predators, Rel. Size, Predators:Habitat, Predators:Rel. Spatial Grain, Predators:Rel. Size | 10 | -1911.88 | 3843.77 | 3.96 | 0.07 |

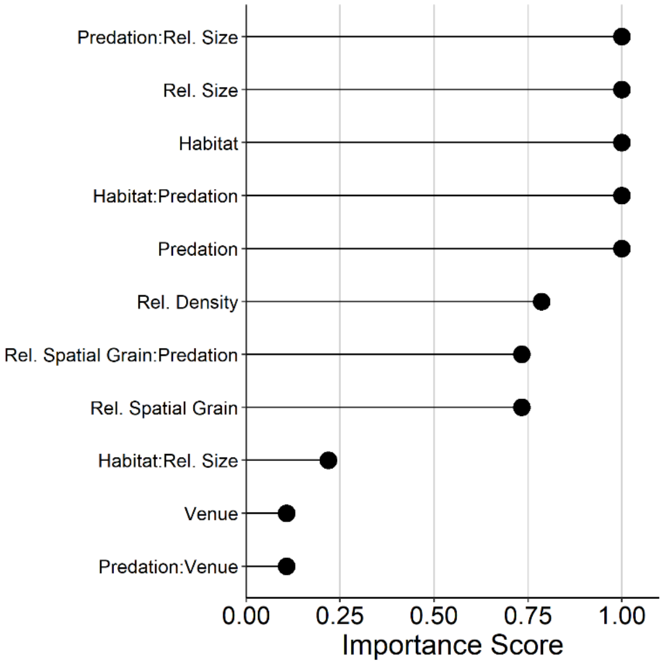


Supplemental Figure 1. Moderator importance scores from models with ΔAIC < 4, as these models have at least some empirical support. Importance scores support the inclusion of terms found in the best model, as all terms included in the best model have importance scores greater than 0.5, which indicates that these predictors were found in models with greater than 0.5 probability of being the best model.

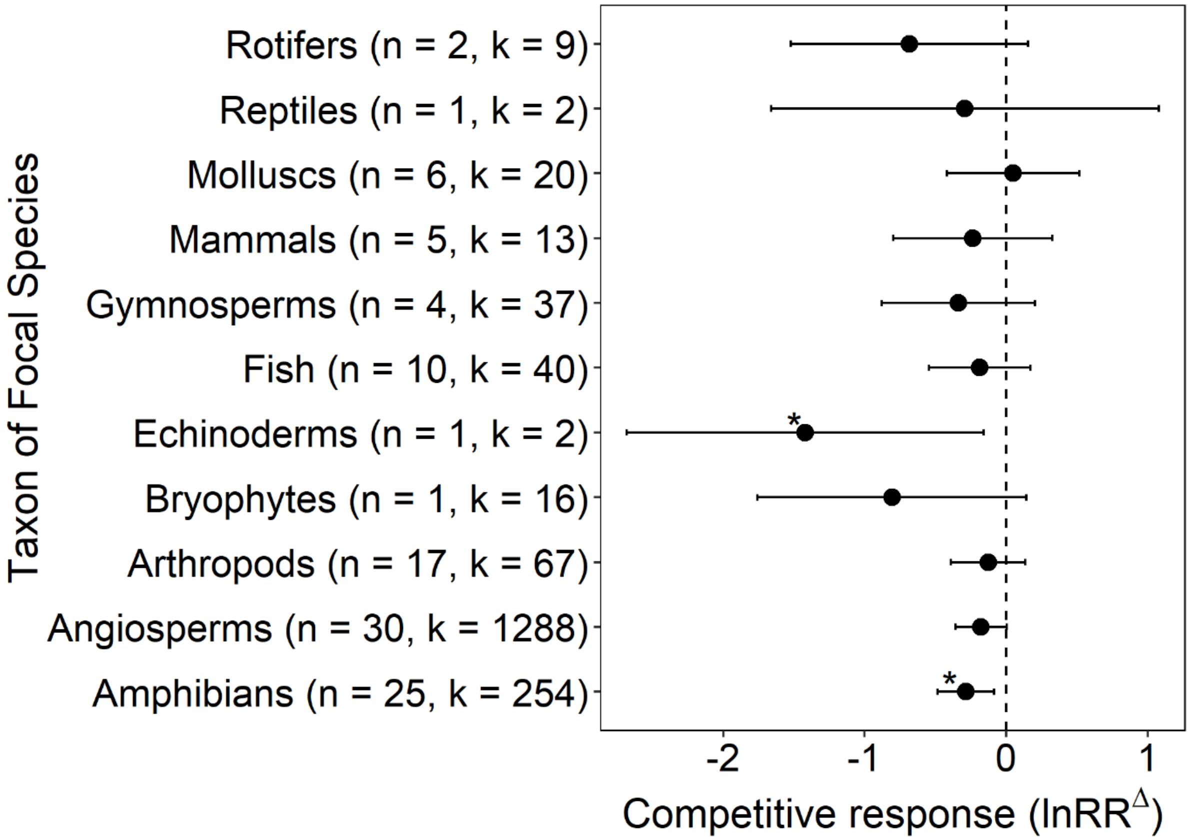


Supplemental Figure 2. Forest plot of the mean effect of focal species taxa on competitive response. Points represent the pooled effect for the taxonomic group and error bars represent 95% confidence intervals generated from a mixed-effects meta-analysis of experiments relating competitive response to the taxonomic group of the focal species. These models included a moderator of taxonomic group, a random effect of study, and the variance-covariance matrix described in the main text. Numbers in the parentheses represent the number of studies (n) and effect sizes (k), respectively, within each taxonomic grouping. The mixed-effects meta-analysis indicated that competitions strength did not vary among taxonomic groups. Asterisks indicate a significant difference from 0 (* $p < 0.05$).

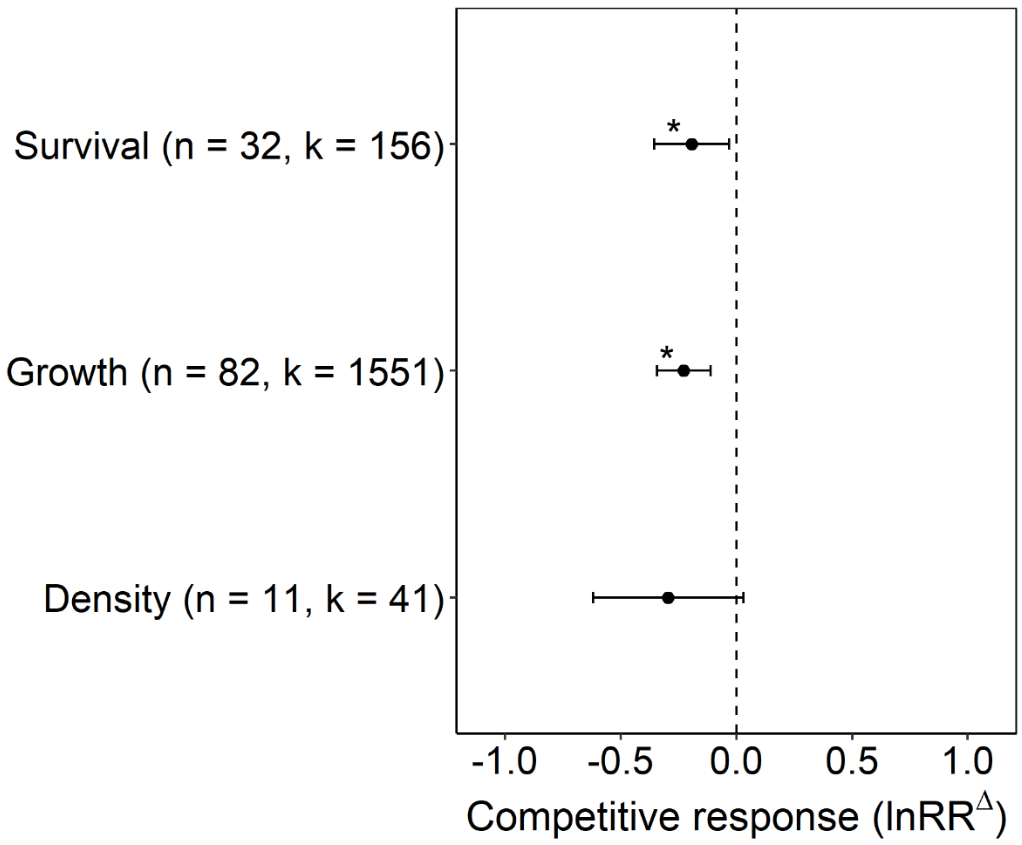


Supplemental Figure 3. Forest plot of the mean effect of study endpoint on competition strength. Points represent the pooled effect for the study endpoint and error bars represent 95% confidence intervals generated from a mixed-effects meta-analysis of experiments relating competition strength to the study endpoint. These models included a moderator of study endpoint, a random effect of study, and the variance-covariance matrix described in the main text. Numbers in the parentheses represent the number of studies (n) and effect sizes (k), respectively, within each study endpoint. The mixed-effects meta-analysis indicated that competition strength did not vary among study endpoint. Asterisks indicate a significant difference from 0 (* $p < 0.05$).

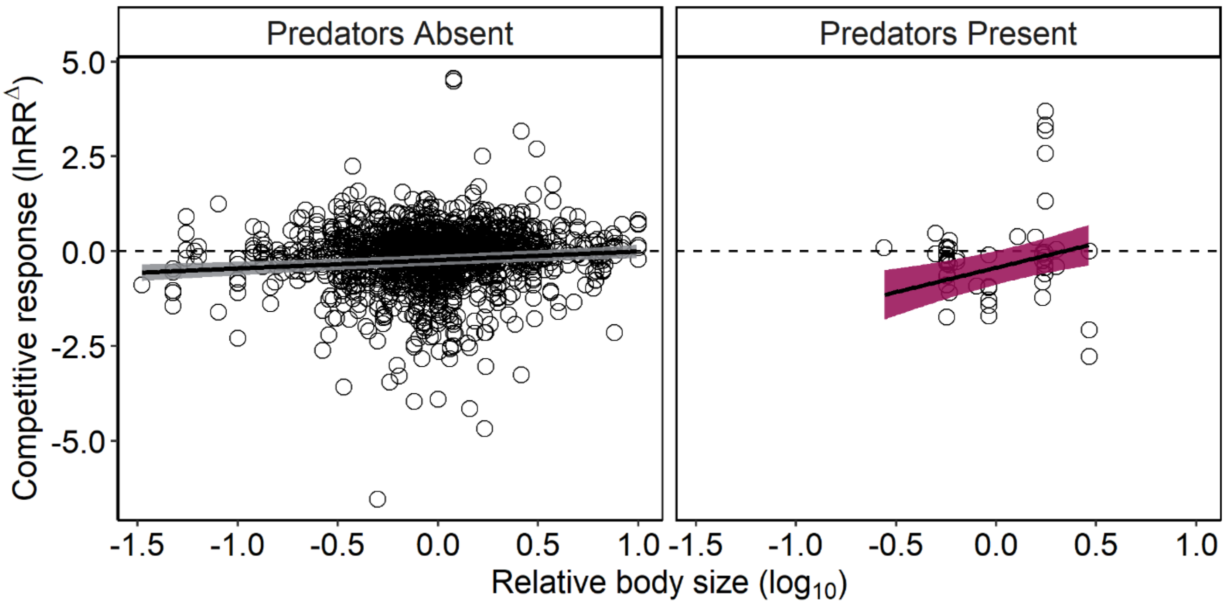


Supplemental Figure 4. Marginal effects plot showing the interactive effects of predation and (A) relative body size (size of focal species (blue) / size of competing species (orange); functional similarity) on competitive response (lnRR$^{\Delta}$). Lines represent marginal effects of relative body size in the mixed-effects meta-regression model and shading shows associated 95% credible bands. To estimate these marginal effects, categorical covariates from the mixed-effects meta-regression model were held at their proportional values and continuous covariates were held at their median value. See Fig. 2 in main text for interpretation of relationship between relative body size and competitive response.

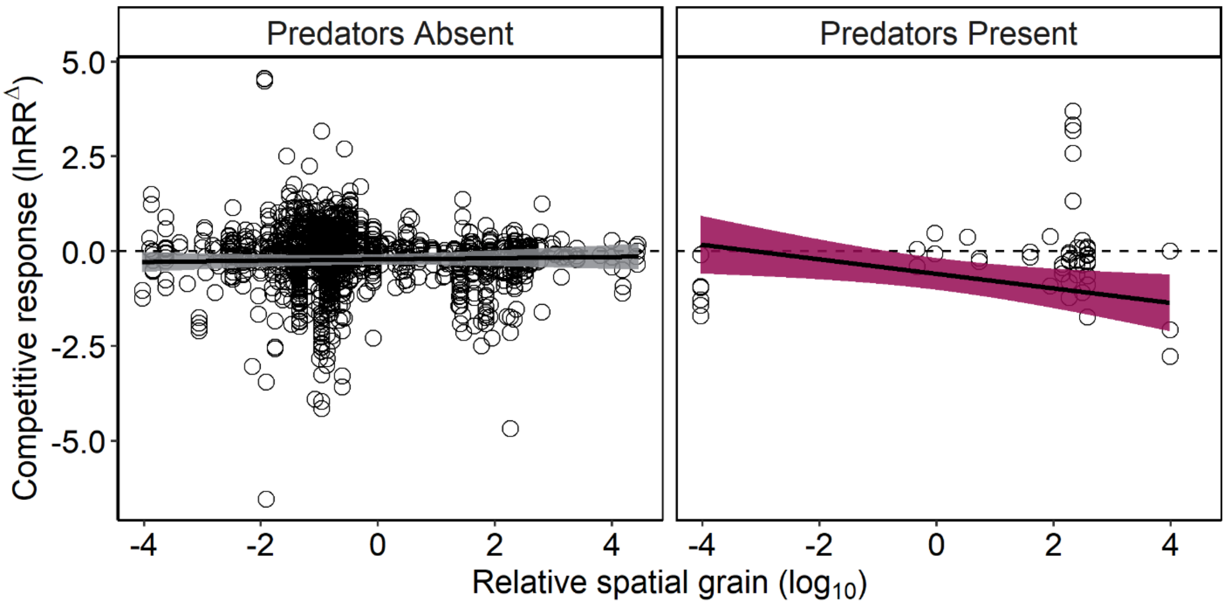


Supplemental Figure 5. Marginal effects plot showing the interactive effects of predation and relative spatial grain (spatial grain / size of focal species) on the outcome of species interactions (lnRR$^{\Delta}$). Lines represent marginal effects of relative spatial grain in the mixed-effects meta-regression model and shading shows associated 95% credible bands. To estimate these marginal effects, categorical covariates from the mixed-effects meta-regression model were held at their proportional values and continuous covariates were held at their median value. See Fig. 2 in main text for interpretation of relationship between relative spatial grain and competitive response.

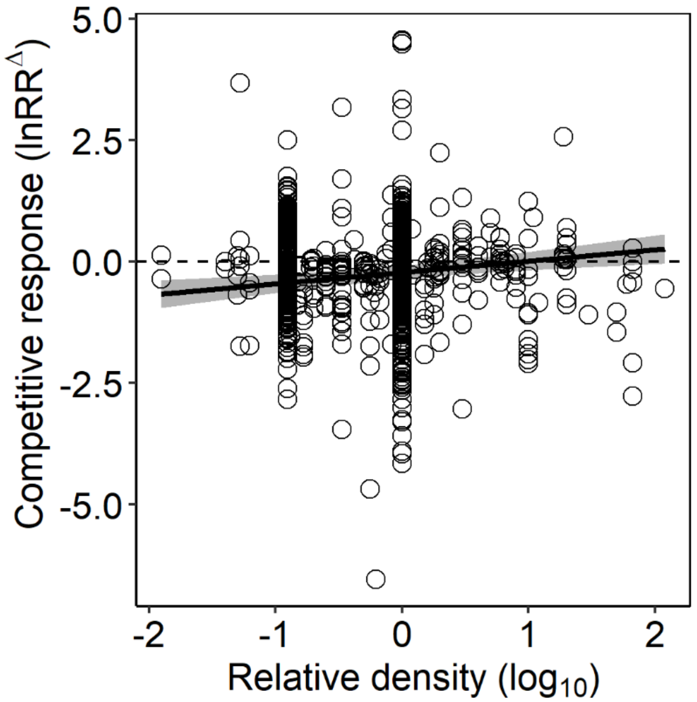


Supplemental Figure 6. Marginal effects plot showing the effects of relative density (density of focal species (blue)/ density of competing species (orange)) competitive response (lnRR$^{\Delta}$). Lines represent marginal effects of relative density included in the mixed-effects meta-regression model and shading shows associated 95% credible bands. To estimate these marginal effects, categorical covariates from the mixed-effects meta-regression model were held at their proportional values and continuous covariates were held at their median value. See Fig. 4 in main text for interpretation of relationship between relative density and competitive response.

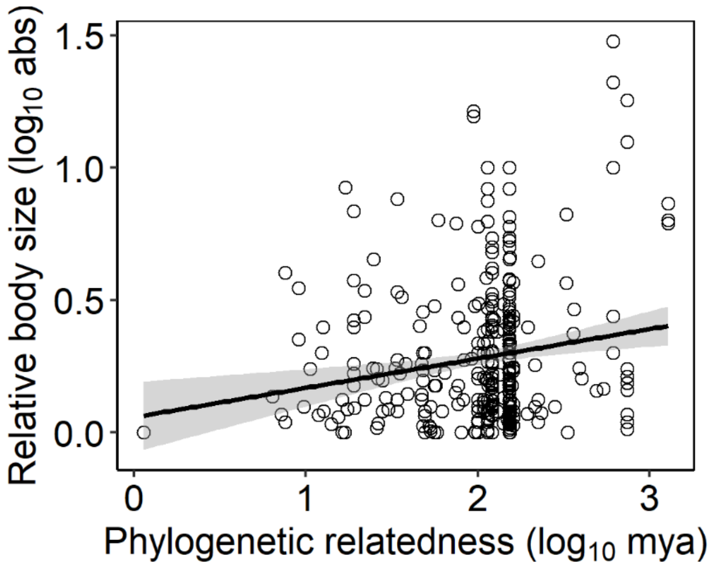


Supplemental Figure 7. Relationship between phylogenetic relatedness (log10 mya) and relative body size (absolute value of log10 size of focal species relative to size of competing species; functional similarity). As species pairs become less phylogenetically related, relative body size differences significantly increased (simple linear regression; $F_{1,422} = 11.59$, $p < 0.001$, $R^2 = 0.03$). This suggests that while our measure of functional similarity exhibited a significant phylogenetic signal, phylogenetic relatedness has little predictive ability on functional similarity.

Supplemental Table 2: Results of a mixed-effects meta-regression model from the model selection indicated best model (lowest AIC) relating competition strength to factors including the phylogenetic relatedness of the competing species, relative density of the focal species, presence of predators, and habitat. Variance estimates of the within and across study random effects are 0.397 and 0.248, respectively. Here, we replaced relative body size from the model in Table 1 with phylogenetic relatedness to test whether phylogenetic relatedness similarly explained competitive responses.

| Moderators | Coefficient | 95% CI | Z-value | P-value |
|---|---|---|---|---|
| Grand Mean | -0.411 | 0.34 | -2.368 | **0.018** |
| Phylogenetic Relatedness (log10) † | 0.084 | 0.13 | 1.274 | 0.203 |
| Relative Density (log10) † | 0.238 | 0.13 | 3.549 | <**0.001** |
| Relative Spatial Grain (log10) † | -0.016 | 0.06 | -0.516 | 0.606 |
| Predators Present | 1.707 | 1.63 | 2.053 | **0.040** |
| Predators Present * Phylogenetic Relatedness | -0.408 | 0.74 | -1.087 | 0.277 |
| Predators Present * Relative Spatial Grain | -0.153 | 0.15 | -1.974 | **0.048** |
| Terrestrial Habitat | 0.002 | 0.26 | 0.013 | 0.990 |
| Predators Present * Terrestrial Habitat | -1.413 | 0.69 | -4.004 | <**0.001** |

Notes: The average outcome of competition is indicated by the grand mean, as the coefficient represents the pooled outcome of competition. For continuous moderators (†), bolded p-values indicate a slope that deviates significantly from zero and the sign of the coefficient indicates the direction of the effect. For categorical moderators, bolded p-values indicate a significant difference from the grand mean with the coefficient indicating the direction and the estimated mean of the category. For interactions between categorical and continuous moderators, bolded p-values indicate a slope that deviates significantly from the continuous moderator and the sign of the coefficient indicates the direction of the effect.

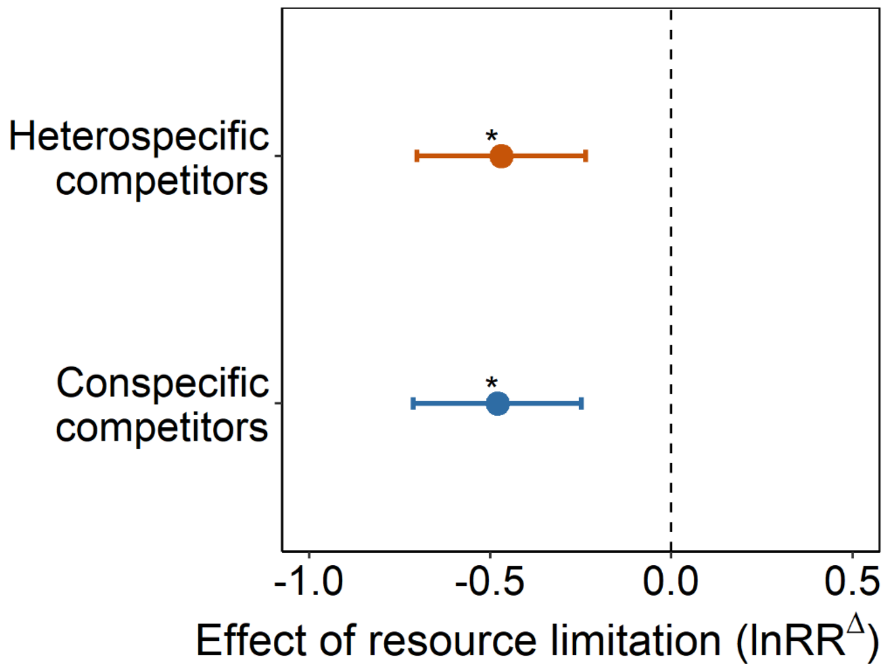


Supplemental Figure 8. Forest plot of the mean effect of resource limitation on various study endpoints when focal individuals are with heterospecifics and conspecifics. Points represent the pooled effect when competitors are heterospecifics and conspecifics and error bars represent 95% confidence intervals generated from a mixed-effects meta-analysis of experiments relating effects of resource limitation to when competitors are heterospecifics and conspecifics. These models included a moderator of heterospecific/conspecific and a random effect of study. The response of these models was the log response ratio of lower resources to higher resources (High → Ambient, High → Low, Ambient → Low). Asterisks indicate a significant difference from 0 (* $p < 0.05$). The mixed-effects meta-analysis indicated that the effect of resource limitation was consistent regardless of whether competitors are heterospecifics or conspecifics (effect of resource limitation on intra- vs. inter-specific competition, $Z = -0.01$, $p = 0.88$). Thus, the consistent negative effects of resource limitation on study endpoints do not translate to changes in our measure of competition strength.
